# Production of homozygous deletion mutants targeting fertilization regulator genes through multiplex genome editing

**DOI:** 10.1101/2025.02.28.640930

**Authors:** Ari Yoshimura, Yura Seo, Sora Kobayashi, Tomoko Igawa

## Abstract

Producing the gene-knockout mutant is a critical strategy in reverse genetics for gene functional analyses. Plant science has applied various mutagenesis, including chemical mutagen, T-DNA insertion, and genome editing. The intact gene expression is completely disrupted, while the mutant retains the intact sequence region. Lately, effective whole open reading frame (ORF) deletion through the CRISPR/Cas9 system using multiplex guide RNAs was reported in Arabidopsis, targeting a gene expressed in somatic cells. Here we applied the scheme targeting the reproductive genes *GENERATIVE CELL SPECIFIC 1*(*GCS1*), *GAMETE EXPRESSED 2* (*GEX2*), *DUF679 DOMAIN MEMBRANE PROTEIN 8* (*DMP8*), and *DMP9*, which are fertilization regulators.

Homozygous gene deletion lines were successfully obtained in the T1 generation, with acquisition rates ranging from 8.3% to 30.0%. The rates approximately correlated with the scores predicted by the DeepSpCas9. The analysis of the *GCS1* deletion line (*Δgcs1*) suggested that avoiding the first bolt removal is important for consistent phenotype of the developed inflorescences. Compared to the previously reported mutants, the *GEX2* deletion line (*Δgex2*) showed a difference in seed development phenotype, indicating that the remaining intact gene region in the mutant could unexpectedly influence the function. Simultaneous deletions of *DMP8* and *DMP9*, which are in distinct chromosomes, were also succeeded in the T1 generation. The obtained results showed that all deletions were inheritable. The scheme challenged here demonstrated the effective production of homozygous mutants for the genes, even if the reproductive lethal or recessive.

## INTRODUCTION

Double fertilization is a unique sexual reproduction manner observed in flowering plants. Two sperm cells are delivered by a pollen tube to an embryo sac, where an egg and a central cell are located. After being released from a pollen tube, one of the sperm cells fertilizes with an egg cell, and another fertilizes with the central cell. The precise progress of two independent fertilization events is directly controlled by protein molecules on the gamete surfaces.

The GENERATIVE CELL SPECIFIC 1 (GCS1), which is also named HAPLESS 2 (HAP2), is the first identified fertilization regulator protein resident specifically in the sperm cell membrane (Mori et al. 2006; vonBesser et al. 2006). Independent studies using distinct T-DNA insertion mutants *gcs1* or *hap2-1* have shown similar insights that GCS1 is an essential factor for fertilization and the bona fide fusogen in plasma membrane fusion with female gametes. However, studies on the functional region required for fertilization have reported different results (Mori Hirai et al. 2010; Wong et al. 2010), remaining controversial. The GAMETE EXPRESSED 2 (GEX2) is a different sperm cell membrane protein that is involved in attachment to the female gametes (Mori et al. 2014). The previous study used two mutant lines, *gex2-1* with point mutation caused by EMS and *gex2-2* with T-DNA insertion. Both mutant lines similarly showed gamete attachment defective and single fertilization phenotypes, whereas different frequencies in seed development phenotypes were observed. The DUF679 DOMAIN MEMBRANE PROTEIN 9 (DMP9) localizes on the sperm cell plasma membrane and contributes more to the egg cell fertilization rather than that of the central cell (Takahashi et al. 2018; Cyprys et al. 2019). The highly similar paralogue DMP8 is known to function redundantly with DMP9, reflected by analyses with RNAi, T-DNA insertion, and genome-edited mutant lines. It has been suggested that DMP8 and DMP9 are also involved in the control of GCS1 distribution in sperm cells (Wang et al. 2022), while the contradict results have been reported from a different study using a distinct *dmp8&dmp9* mutant genotype (Shiba et al. 2023).

All the mutagenesis by T-DNA insertion, chemical treatment, and CRISPR/Cas9 system using single guide RNA, remain partially intact sequences of the original gene in the mutant genome. The possibility of unexpected interference due to abnormal transcripts and proteins derived from the remaining sequences cannot be eliminated. Regarding molecular functions of the fertilization regulators, the inconsistency reported from different studies could be due to the use of distinct genotype mutant lines in each study. Recently, effective deletion of larger chromosomal segments by CRISPR/Cas9 system has been reported in Arabidopsis (Ordon et al. 2023). Complete elimination of the ORF can avoid potential interference in expression. Therefore, we applied this strategy to produce *GCS1, GEX2, DMP8*, and *DMP9* mutant lines, in which the whole ORF region is deleted from the genome. The homozygous ORF deleted lines were also obtained in T1 generations at substantial frequency, allowing the analyses with homozygous mutants that cannot be obtained due to lethal or uninheritable traits. Compared with the mutants used in previous studies, we show the importance of ORF deletion for precise gene functional analysis.

## RESULTS

### Construction of the CRISPR/Cas9 vectors for gene deletion

The *zCas9i* is a *Zea mays* codon-optimized and intronized *Cas9* coding sequence, with nuclear localization signals (NLS) at both terminal ends. The zCas9i has greatly improved the effect of Arabidopsis genome editing (Grützner et al. 2020). Furthermore, the use of two pairs of guide RNAs (four guide RNAs) sandwiching the deletion target region was best for chromosomal deletion (gene deletion) (Ordon et al. 2023). Utilizing these schemes and target sequences selected based on DeepSpCas9 prediction (Kim et al. 2019), vectors for each fertilization regulator gene deletion were constructed (Figure S1). For *GCS1*, target sequences at the 3′ side were designed within ORF to avoid influencing the neighbor gene expression (Fig. 1a).

**Fig. 1.**
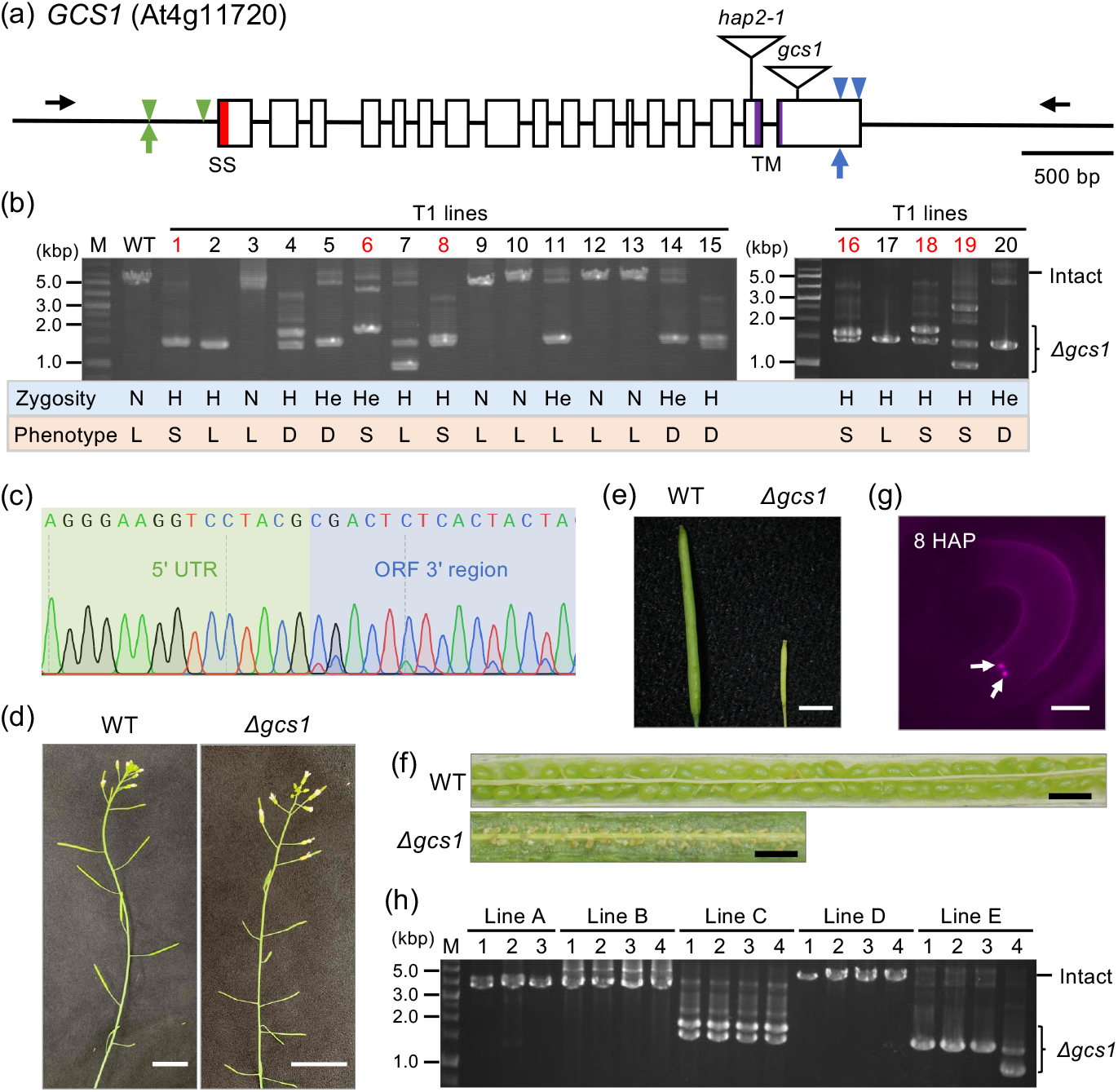
Evaluation of *GCS1* deletion. **(a)** Schematic drawing of genomic *GCS1*, which is comprised of 17 exons (white boxes) and 16 introns (internal lines). Green and blue triangles represent target sites at 5′ upstream and 3′ downstream of the ORF. Black arrows are primers used for genomic PCR to check the ORF deletion. Open triangles with a vertical line indicate the T-DNA insertion sites in *hap2-1* and *gcs1* mutants (vonBessor et al., 2006; Mori et al. 2006). The corresponding signal sequence (SS) and transmembrane (TM) regions are red and purple colored, respectively. **(b)** Genomic PCR with primers shown in (a) to check the gene deletion in T1 transgenic lines. The positions of intact *GCS1* (ca. 5.2 kbp) and the gene deletion (*Δgcs1*) (ca. 1.0–2.0 kbp) amplicons are marked with a line and a curly bracket, respectively. The estimated zygosities of gene deletion based on the amplicon patterns are represented below the photo (H; homozygous, He; heterozygous, N; no deletion). The letters at the bottom indicate the silique phenotype on the inflorescence observed in each T1 plant (L; long silique, S; short silique, D; both long and short siliques). M; size marker. **(c)** Sequencing result of a band from line 1 showing *GCS1* homozygous deletion (*Δgcs1*). Green and blue arrows in (a) are the junction of 5′ UTR and OFR 3′ region. **(d)** Inflorescences of wild type (WT) and *Δgcs1*. bar = 2 cm. **(e)** WT and *Δgcs1* siliques. bar = 3 mm. **(f)** Seed development in WT and *Δgcs1* siliques. bar = 1 mm. **(g)** Double fertilization block caused by *Δgcs1* sperm cells. Two sperm nuclei (white arrows) are arrested in an embryo sac. HAP; hours after pollination. bar = 40 µm. **(h)** Genomic PCR performed as described in (a) to check the deletion patterns in different inflorescences (1–4) in each transgenic line (A-E). The amplicons marked with a line and a curly bracket reflect the intact *GCS1* amplicon and smaller bands suggestive of gene deletion. M; size marker.

### Gene deletion targeting *GCS1*

*GCS1* deletion in the obtained T1 transgenic plant lines was evaluated by genomic PCR with a primer pair annealing to the outside of the deletion target region (Fig. 1a). The genome DNA isolated from a cauline leaf on the first bolted inflorescence was used for PCR. The larger amplicon (5202 bp) reflects intact *GCS1*, and the smaller bands (ca. 1.4–1.8 kbp) indicate the gene deletion. Among 20 transgenic lines examined, 15 lines indicated gene deletion amplicon patterns (75.0% deletion efficiency) (Fig. 1b). In the prescreening by PCR, 10 lines are estimated as the homozygous deletion genotype, as only the smaller bands were detected. Sequencing of the smaller band detected in line 1 revealed the deletion of the *GCS1* coding region (Fig. 1c). Therefore, lines showing only smaller bands were suggestive of the homozygous *GCS1* deletion genotypes (30.0% acquisition rate). The genomic PCR evaluation also detected multiple larger and smaller bands in most of the lines (Fig. 1b). This was due to the different deletion patterns, such as deletions occurring between the 5′ or 3′ target sites and deletions of various lengths in each chromosome locus, as the mixed waveforms from around the target site were detected in sequencing (data not shown).

The first bolted inflorescence was pinched in these transgenic lines to stimulate the development of lateral inflorescences. Finally, six transgenic lines (labeled in red in Fig. 1b) exhibited short siliques containing undeveloped seeds in all emerged inflorescences (Fig. 1d, e, f). The sperm cells from these lines could not fuse with the female gametes (Fig. 1g), confirming these were the homozygous *gcs1* deleted plants (*Δgcs1*). A heterozygous *Δgcs1* (+/*Δgcs1*, from the line 1 in Fig. 1b) was produced by crossing with HITR10mRFP as the male plant. The normal seed development frequency in +/*Δgcs1* was similar to that of +/*gcs1*, as previously reported (Figure S2).

### Gene deletion pattern in an inflorescence is consistent in T1 plants

In the present evaluation, six plants exhibiting homozygous *gcs1*-knockout phenotype were obtained in the T1 generation, resulting in a 30% acquisition rate. All the lines estimated as deletion negative in the genomic PCR were fertile (lines 3, 9, 10, 12, and 13). Almost all lines suggested as homozygous deletions were infertile, although line 2 showed an exception. The overall consistency between the estimation by genomic PCR and the phenotypes was 60% (Fig. 1b). Additionally, five lines (lines 4, 5, 14, 15, and 17) produced inflorescences that contained either long siliques with developed seeds or short siliques with undeveloped seeds. The silique phenotype was consistent within each inflorescence. The zCas9i is driven by a constitutive Arabidopsis *RPS5A* promoter, which is also active in the meristem (Tsutsui & Higashiyama 2017). Therefore, these mixed phenotypes observed may indicate that different deletion patterns occurred independently in each inflorescence meristem.

We then examined whether the inflorescences within the same line exhibited different deletion patterns and phenotypes under the condition that the first inflorescence was not pinched. We produced five additional transgenic T1 lines were produced, and three to four inflorescences from each line were analyzed by genomic PCR (Fig. 1h). The results showed that almost all the inflorescences within the same line displayed identical PCR patterns, except for one inflorescence in line E, which produced an additional smaller band. Among the five lines, line C exhibited two smaller bands but not the *GCS1* intact amplicon, and all the inflorescences produced short siliques with undeveloped seeds, indicating complete infertility.

Refraining from removing the first inflorescence led to a more consistent phenotype among the developing inflorescences, which was guaranteed by the same deletion pattern. Considering these factors, this approach enabled the production of a homozygous mutant line in the T1 generation, even when a defect in the target gene is lethal to reproduction.

### Gene deletion targeting *GEX2*

The sperm cell-expressed protein GEX2 plays a significant role in the attachment to female gametes (Mori et al. 2014). GEX2 is not reproductive essential, thus homozygous *gex2* mutants (*gex2-1* and *gex2-2*) can be maintained and were analyzed in a previous study. Due to attachment failures, double fertilization block and single fertilization of the egg or central cell occur in the *gex2* mutants. Therefore, three seed development phenotypes are observed in the *gex2* mutants: undeveloped seeds, which arise from double fertilization failure or single fertilization of the egg cell; aborted seeds, resulting from single fertilization of the central cell; and normal seeds by success of double fertilization (Fig. 2d).

**Fig. 2.**
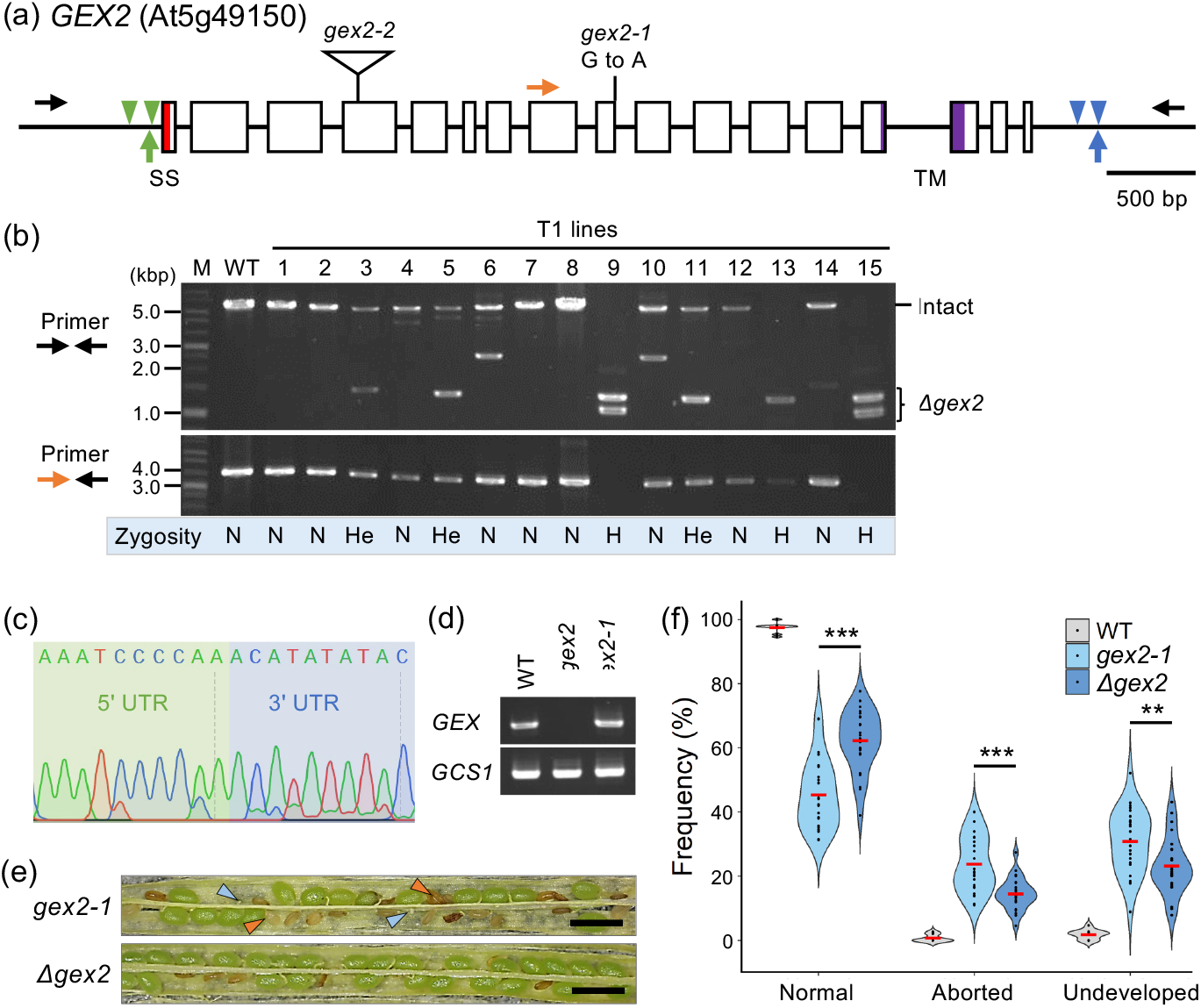
Evaluation of *GEX2* deletion. **(a)** Schematic drawing of genomic *GEX2*, which is comprised of 17 exons (white boxes) and 16 introns (internal lines). Green and blue triangles represent target sites at 5′ upstream and 3′ downstream of the ORF. Black and orange arrows are primers used for genomic PCR to check the deletion. The first G in the ninth intron is changed to A in the *gex2-1* mutant (Mori et al., 2014). An open triangle with a vertical line indicates the T-DNA insertion site in *gex2-2* mutant (Mori et al., 2014). The corresponding signal sequence (SS) and transmembrane (TM) regions are red and purple colored, respectively. **(b)** Genomic PCR evaluation of T1 transgenic lines with primers that anneal outside the ORF (upper photo) or primers that anneal outside and inside the ORF (lower photo). The arrows at the left of each photo are the used primer sets. The intact *GEX2* amplicon (ca. 6.8 kbp) and the smaller bands suggesting deletions (ca. 1.0–1.5 kbp) are marked with an arrowhead and a curly bracket, respectively. The estimated zygosities of gene deletion based on the amplicon patterns are represented below the photo (H; homozygous, He; heterozygous, N; no deletion). M; size marker. **(c)** Sequencing result of a band from line 13 (upper photo in (b)) showing *GEX2* homozygous deletion (*Δgex2*). **(d)** RT-PCR for partial *GEX2* in WT, *gex2-1*, and *Δgex2* (upper photo). *GCS1* was amplified as the internal control (lower photo). **(c)** Siliques from *gex2-1* (upper) and *Δgex2* (lower). Pale blue and orange triangles indicate aborted and undeveloped seeds. bar = 1 mm. **(d)** Violin plots of frequencies of each seed phenotype observed in WT (gray), *gex2-1* (pale blue), and *Δgex2* (smoke blue) siliques. Twelve, 26, and 25 siliques from WT, *gex2-1*, and D*gex2* were dissected for seed counting. The black dots and a red horizontal bar in each plot represent each frequency per silique and the median. Significant differences between *gex2-1* and *Δgex2* detected with Welch’s t-test are indicated with asterisks (***; *p* < 0.001, **; *p* < 0.01).

Gene deletion for *GEX2* was performed, and fifteen T1 plants were analyzed by genomic PCR (Fig. 2). With the primers that annealed outside the *GEX2* ORF (Fig. 2a), six lines displayed the deletion (40.0% efficiency). Moreover, three of them were estimated as homozygous deletion lines (lines 9, 13, and 15 in Fig. 2b), showing only smaller bands (ca. 1.0–1.5 kb) but not intact *GEX2* (6838 bp). Additional genomic PCR with primers that annealed outside and inside the ORF did not produce the amplicon, also supporting homozygous deletion. Sequencing of the smaller band of line 13 revealed the complete *GEX2* deletion (*Δgex2*) (Fig. 2c), resulting in a 20.0% acquisition rate in the T1 generation.

### The *Δgex2* exhibited higher fertility than the *gex2-1*

Plants that retain *Δgex2* loci but not T-DNA were selected from the selfed progeny and were used for successive analysis. Similar to the phenotypes reported in the *gex2-1* and *gex2-2* mutants, three seed development phenotypes were observed in the *Δgex2* plant (Fig. 2d). In comparison with the *gex2-1* mutant, each seed phenotype frequency was significantly different, particularly normal seed frequency was higher in the *Δgex2* (Fig. 2e). Although the *gex2-2* was not analyzed in this study, the frequencies observed in the *Δgex2* were similar to those reported for *gex2-2* (Mori et al. 2014). The *gex2-1* has a single nucleotide substitution that disrupts the splicing of the 9th intron, resulting in a premature termination codon (Fig. 2a). RT-PCR analysis revealed the presence of partial *GEX2* transcripts in the *gex2-1* but not in the *Δgex2* (Fig. 2f).

### Gene deletion targeting *DMP8, DMP9*, and *DMP8*&*DMP9*

Among the 10 proteins in the Arabidopsis DMP family, DMP8 and DMP9 are specific to sperm cell plasma membrane and play a significant role in egg cell fertilization in double fertilization (Takahashi et al. 2018; Cyprys et al. 2019). While the *dmp8* null mutant does not exhibit any abnormal phenotype, it is known that DMP8 functions redundantly with DMP9. The *dmp9* mutant shows fertilization failure, and the phenotype is enhanced in the *dmp8*&*dmp9* double mutant (Cyprys et al. 2019; Wang et al. 2022; Shiba et al. 2023).

In the genomic PCR analysis of the twenty-four T1 transgenic plants that introduced CRISPR/Cas9 targeting *DMP8* deletion, 13 lines displayed smaller bands (ca. 1.3–1.5 kbp), resulting in 54.2% deletion efficiency (Fig. S3). Among these, two lines (lines 21 and 23) exhibited deletion-homozygous patterns (Fig. S3b). Sequencing of the band from line 21 revealed a complete *DMP8* ORF deletion, resulting in an 8.3% acquisition rate (Fig. S3c). For *DMP9*, the smaller deletion bands (ca. 1.3–1.7 kbp) were detected in 14 lines among 22 transgenic T1 lines, showing 63.6% deletion efficiency (Fig. 3a, b). Four lines (lines 2, 3, 9, 22) among these displayed smaller bands but not intact *DMP9* amplicon, indicating homozygous deletion. Sequencing of the band from line 3 indicated a complete *DMP9* deletion (*Δdmp9*); thus, the resulting acquisition rate was 18.2%.

**Fig. 3.**
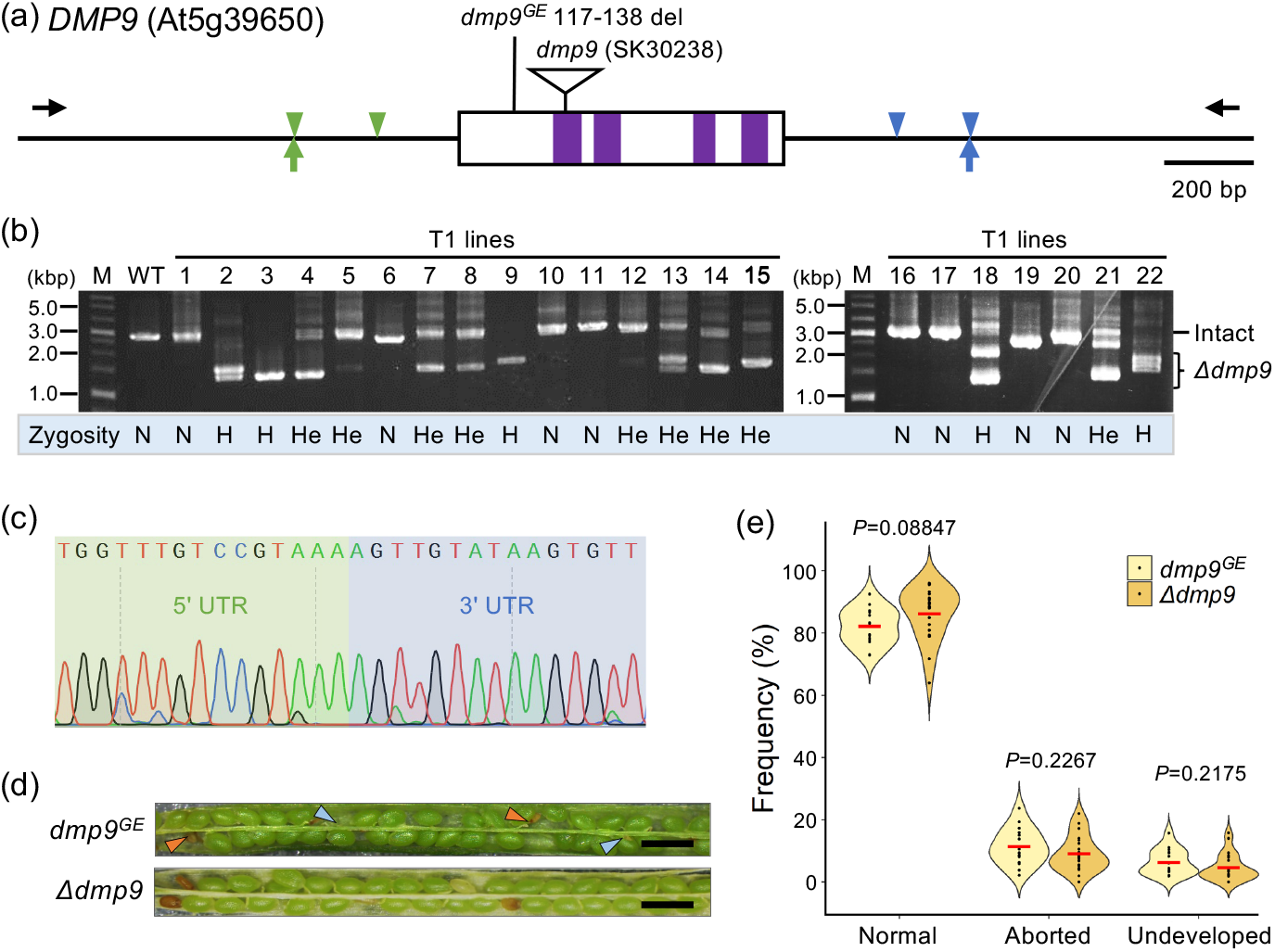
Evaluation of *DMP9* deletion. **(a)** Schematic drawing of genomic *DMP9*, an intronless gene. Green and blue triangles represent target sites at 5′ upstream and 3′ downstream of the ORF. Black arrows are primers used for genomic PCR to check the deletion. A vertical line is a genome-edited site (117-138 deletion) in the *dmp9*^*GE*^ line (Shiba and Takahashi et al. 2023) and an open triangle with a vertical line indicates the T-DNA insertion site in *dmp9* (SK30238) (Wang et al. 2022). Four transmembrane regions are colored in purple. **(b)** Genomic PCR evaluation of T1 transgenic lines with primers that anneal outside the ORF (black arrows in (a)). The intact *DMP9* amplicon (ca. 2.9 kbp) and the smaller bands (ca. 1.0–1.7 kbp) suggesting deletions are marked with a line and a curly bracket, respectively. The estimated zygosities of gene deletion based on the amplicon patterns are represented below the photo (H; homozygous, He; heterozygous, N; no deletion). M; size marker. **(c)** Sequencing result of the smallest band from line 21 indicating *DMP9* homozygous deletion (*Δdmp9*). **(d)** Violin plots of frequencies of each seed phenotype observed in *dmp*^*GE*^ (pale yellow) and *Δdmp9* (light orange) siliques. Fifteen and 24 siliques from *dmp*^*GE*^ and *Δdmp9* were dissected for seed counting. The black dots and a red horizontal bar in each plot represent each frequency per silique and the median. P-values between *gex2-1* and *Δgex2* detected with Welch’s t-test are indicated above the plots.

In the present study, the simultaneous deletion of two OFRs, *DMP8* and *DMP9*, was challenged. Four sets of guide RNAs (eight target sites in total) expression cassettes were constructed in the T-DNA region (Fig. S1b). The *DMP8* and *DMP9* deletions were evaluated in 33 transgenic T1 lines by genomic PCR. The *DMP8* deletion occurred in five lines, and none of them suggested homozygous deletion. For *DMP9*, deletion occurred in 13 lines, and four lines among them were estimated as deletion homozygous. In this evaluation, lines exhibited homozygous deletion of both genes were not obtained. Alternatively, two of the *DMP8* heterozygous and the *DMP9* homozygous deletion lines (lines 9 and 22) were obtained at 6.1% efficiency (Fig. S4f). The *Δdmp8*&*Δdmp9* plant without T-DNA transgene was established from the selfed progeny.

### The *Δdmp9* showed a similar phenotype with *dmp9*^*GE*^

DMP9 is a significant player in egg cell fertilization, so previously reported *dmp9* mutants exhibited aborted seed phenotype caused by single fertilization of the central cell, in addition to undeveloped seed due to double fertilization failure (Fig. 3d). The *Δdmp9* plant without T-DNA transgene was selected from the selfed progeny. The seed development phenotype was evaluated in *Δdmp9* compared to a genome-edited mutant *dmp9*^*GE*^ (Shiba and Takahashi et al. 2023). As a result, no significant differences in each seed phenotype between the *Δdmp9* and *dmp9*^*GE*^ were detected (Fig. 3e).

## DISCUSSION

### Gene deletion efficiency correlated with DeepSpCas9 score

Successful gene deletion by multiplex genome editing has been reported targeting Arabidopsis *WRKY30*, coding a transcription factor expressed in somatic cells (Ordon et al. 2023). We applied this scheme for gene deletions of male gamete-expressed genes *GCS1, GEX2, DMP8*, and *DMP9*, which encode players in double fertilization control. The deletion efficiency was approximately correlated with the DeepSpCas9 scores (Fig. S1a). Using target sequences with more than 50% score for all four guide RNAs especially led to higher deletion efficiency. Although some target sites had lower scores, homozygous deletion lines for each gene were successfully obtained in the T1 generations, with an acquisition rate ranging from 8.3% to 30.0%. The doubly deleted *Δdmp8*&*Δdmp9* lines were not found in the 33 transgenic lines in the T1 generation (Fig. S4). This absence may be attributed to the low DeepSpCas9 score associated with the *DMP8* target sequences. The predicted efficiency by the DeepSpCas9 score for each gene could have a redundant effect on the deletion of multiple genes. The *dmp8* heterozygous and *dmp9* homozygous deletions were achieved in the T1 generation (2/33, 6.1%) (Fig. S4f), and the *Δdmp8*&*Δdmp9* plants were obtained in the selfed progeny, also showing the deletions were inheritable, as well as *Δgcs1* (Fig. S2) and *Δgex2* (Fig. 2f).

The production of homozygous *gcs1*^-/-^ individuals has been reported by Nagahara et al. (Nagahara et al. 2015). The homozygous gcs1 were rescued by introducing GCS1, which can later be removed in the mature progeny seed stage using a heat-inducible Cre-*loxP* recombination system. In contrast to previous methods, the present study produced *GCS1* homozygous deletion lines in the T1 generation. The gene deletion scheme applied here has a great advantage, especially for functional analysis of reproductive essential and recessive genes.

### Genome editing pattern is consistent in each inflorescence

Genome editing occurs independently in each transgenic cell, resulting in cells with different editing patterns co-existing within the T1 plant. Nevertheless, the silique phenotype was consistent within an inflorescence in the lines targeted *GCS1* deletion. The RPS5A promoter is highly active in the meristem (Weijers et al. 2001), suggesting that genome editing occurred at an early stage of inflorescence development, thus the inflorescences differentiating from the edited cell exhibited the same silique phenotype. After the first bolted inflorescence was pinched, independent genome editing occurred in the lateral inflorescence meristem. Consequently, some meristematic cells may have escaped gene deletion, and the developed inflorescence exhibited long silique with developed seeds, resulting from double fertilization. Refraining from pinching the first inflorescence maintains apical dominance. The result that all inflorescences from the same T1 line displayed nearly identical genome editing patterns, provided that apical dominance remained undisturbed, indicates that the developed inflorescences were further differentiated from the meristem of the main inflorescence. The speculation would be clarified by cytological analyses of inflorescence meristem differentiation under disturbed or maintained apical dominance. In the present study, avoiding the removal of the first inflorescence effectively results in uniform phenotypes caused by the same genome editing patterns, even if the total number of inflorescences is reduced.

### Gene deletion is a potential tool for precise molecular functional analyses

In comparison with the gene deletion lines (*Δgcs1, Δgex2, Δdmp9*) and the T-DNA insertion and genome-edited mutant lines, no significant differences were detected in each phenotype and the degree, except for *gex2-1*, a mutant with a single nucleotide substitution. Although the difference in frequencies of the phenotypes between *gex2-1* and ge*x2-2* was not focused on the previous study (Mori et al. 2014), our result supported the finding. The fact of the partial *GEX2* transcript accumulation (Fig. 2c) suggested the potential interference on the phenotype observed in the *gex2-1* mutant, although the underlined phenomenon was not investigated. To avoid such unexpected artifacts completely, gene deletion is a powerful tool for precise functional analyses. Complement analyses, which introduce the intact or variant gene into a knockout mutant, are a major strategy for investigating molecular functional regions. The GCS1 protein comprises a long ectodomain and a short endodomain via a single transmembrane region (Fig. 1a). As of today, the contribution of the endodomain in gamete fusion remains controversial. Independent studies have reported inconsistent results using *hap2-1* or *gcs1* mutants, where T-DNA is inserted in the upstream or the downstream proximity of a transmembrane region (Fig. 1a). The difference in the disturbed position of the GCS1 could have influenced the protein function. Therefore, *Δgcs*1 created in this study would be a good background line for precise functional analyses, which are ongoing in our laboratory.

## MATERIALS AND METHODS

### Plant growth conditions and transformation

The sperm cell nuclear marker line, HTR10mRFP (*Arabidopsis thaliana* Col-0 background) (Ingouff et al. 2007), was used to produce gene deletion lines. Plants were grown at 22°C under the 16 h light / 8 h dark cycle or 25°C under the continuous light condition. Arabidopsis transformation was performed by floral inoculation (Narusaka et al. 2010) using *Agrobacterium tumefaciens* GV3101. The seeds expressing RFP were selected as independent T1 lines (Shimada et al. 2010). In the T2 population of each line, the RFP-negative seeds were selected as the T-DNA-free plants with gene deletion.

### Target sequence selection and plasmid constructions

The target sequences for the CRISPR/Cas9 system were selected within the 5’ and 3’ untranslated regions (UTR), where ca. 300 to 600 bp far from the start and stop codon. As an exception, 3’ target sequences on the ORF were used for GCS1 deletion to not influence the downstream neighboring gene. There was at least a 100 bp gap between the two target sequences at each 5’ and 3’ sides. Finally, two pairs of target sequences with the highest scores detected in DeepSpCas9 (Kim et al. 2019) were chosen for each gene. For *GCS1* deletion, a modified pDGE1108 vector (Ordon et al. 2023), which contains a *NLS-zCas9i-NLS* expression cassette under the controls of Arabidopsis *RPS5A* promoter and the triple terminator (fusions of 35S terminator from Cauliflower Mosaic Virus, *Nicotiana benthamiana Actin3* terminator region, and the *Rb7* matrix attachment region from *N. tabacum*) (Diamos and Mason 2018), and a FAST marker gene cassette (Shimada et al. 2010) was used. A region between the BsaI sites was replaced with the multiple Gateway^®^ cassettes from pDe_Cas9 vector (Fauser et al. 2014), producing a pDGE1108-(GW) vector. Each *GCS1* target sequence was cloned into pMR217 (pENTR_L1R5_AtU6gRNA), pMR219 (pENTR_L5L4_AtU6gRNA), pMR204 (pENTR_R4R3_AtU6gRNA) and pMR205 (pENTR_L3L2_AtU6gRNA) (Ritter et al. 2017; Swinnen et al. 2022), respectively, and then the four guide RNA cassettes were inserted into pDGE1108-(GW) with MultiSite Gateway^®^ system (ThermoFisher Scientific Inc., Tokyo, Japan). For *GEX2, DMP8*, and *DMP9* deletions, four target sequences were separately cloned into pMR217 or pMR205 vectors. The BsaI-cut and ligation reactions were performed as reported (Stuttmann et al. 2021) to insert four cassettes into a pDGE1108 vector. For the double deletions of *DMP8*&*DMP9*, the four guide RNA clusters for each *DMP8* and *DMP9* deletion, which were separately cloned into the pDGE1108 vectors, were ligated to a single pDGE1108. The primers used for plasmid constructions are shown in Table S1.

### Gene deletion check

The primers used for gene deletion check are shown in Table S1. Genomic DNA was isolated from cauline leaves on the first developed inflorescence as described by Ishizaki et al. (Ishizaki et al. 2013). To evaluate deletion patterns between the inflorescences, genomic DNA was isolated from 3-4 inflorescence stems that emerged after cutting the first inflorescence in each transgenic line. The PCR reactions were performed with KOD One^®^ PCR Master Mix (TOYOBO CO., LTD., Osaka, Japan). The PCR amplicons suggestive of gene deletions were excised from the gel, and then the purified fragments were sequenced.

### RT-PCR

Total RNA was extracted from mature flowers using an RNeasy Plant Mini Kit (QIAGEN N. V., Venlo, Nederland). The cDNA was synthesized from 200 ng of total RNA using the ReverTra Ace^®^ qPCR RT Master Mix (TOYOBO CO., LTD.). The *GEX2* was amplified using primers GEX2 ORF5P and GEX2 RT3P, and *GCS1* was amplified as the control gene, using GCS1 RT5P and GCS1 RT3P.

### Evaluation of seed development

After 2 to 3 weeks of the anthesis, the developing siliques were dissected under an SZX9 stereomicroscope (EVIDENT, Ina, Japan). The number of normal, aborted, and undeveloped seeds were counted, and 12-26 siliques were observed for each transgenic line. As the control, HTR10-mRFP plants were used. The images were captured with a digital camera DP72 and cellSence standard software (EVIDENT).

### Observation of fertilization

For observation of fertilization phenotype, the wild type was emasculated on the day before flowering and then pollinated with *Δgcs1* pollen. Ovules at 8 hours after pollination (HAP) were dissected from a pistil and observed under the fluorescent microscope IX81 (EVIDENT). The images were captured with a digital camera DP74 and cellSence software (ver. 4.2) (EVIDENT).

### Statistical analysis

The data were analyzed using Welch’s t-test for each seed development phenotype.

## ACKNOWLEDGMENT

Dr. Masaki Endo (NARO) kindly provided pMR204, pMR205, and pMR217 vectors. Mr. Burbudi B. Pratama and Ms. Tomoe Kano helped with the plant cultivation and seed collection.

## SUPPLEMENTAL INFORMATION

Supplemental information is available at xx Online.

## REFERENCES

Cyprys P, Lindemeier M, Sprunck S (2019) Gamete fusion is facilitated by two sperm cell-expressed DUF679 membrane proteins. Nat Plants 5:253–257

Diamos AG, Mason HS (2018) Chimeric 3’ flanking regions strongly enhance gene expression in plants. Plant Biotechnol J 16:1971–1982

Fauser F, Schiml S, Puchta H (2014) Both CRISPR/Cas-based nucleases and nickases can be used efficiently for genome engineering in Arabidopsis thaliana. Plant J 79:348–359

Grützner R, Martin P, Horn C, Mortensen S, Cram EJ, Lee-Parsons CWT, Stuttmann J, Marillonnet S (2021) High-efficiency genome editing in plants mediated by a Cas9 gene containing multiple

Ingouff M, Hamamura Y, Gourgues M, Higashiyama T, Berger F (2007) Distinct dynamics of HISTONE3 variants between the two fertilization products in plants. Curr Biol 17:1032–1037

Ishizaki K, Johzuka-Hisatomi Y, Ishida S, Iida S, Kohchi T (2013) Homologous recombination-mediated gene targeting in the liverwort Marchantia polymorpha L. Sci Rep 3:1532

Kim HK, Kim Y, Lee S, Min S, Bae JY, Choi JW, Park J, Jung D, Yoon S, Kim HH (2019) SpCas9 activity prediction by DeepSpCas9, a deep learning-based model with high generalization performance. Sci Adv 5:eaax9249

Mori T, Hirai M, Kuroiwa T, Miyagishima SY (2010) The functional domain of GCS1-based gamete fusion resides in the amino terminus in plant and parasite species. PLoS One 5:e15957

Mori T, Igawa T, Tamiya G, Miyagishima S-y, Berger F (2014) Gamete Attachment Requires GEX2 for Successful Fertilization in Arabidopsis. Curr Biol 24:170–175

Mori T, Kuroiwa H, Higashiyama T, Kuroiwa T (2006) GENERATIVE CELL SPECIFIC 1 is essential for angiosperm fertilization. Nat Cell Biol 8:64–71

Nagahara S, Takeuchi H, Higashiyama T (2015) Generation of a homozygous fertilization-defective gcs1 mutant by heat-inducible removal of a rescue gene. Plant Reprod 28:33–46

Narusaka M, Shiraishi T, Iwabuchi M, Narusaka Y (2010) The floral inoculating protocol: a simplified Arabidopsis thaliana transformation method modified from floral dipping. Plant Biotech 27:349–351

Ordon J, Kiel N, Becker D, Kretschmer C, Schulze-Lefert P, Stuttmann J (2023) Targeted gene deletion with SpCas9 and multiple guide RNAs in Arabidopsis thaliana: four are better than two. Plant Methods 19:30

Ritter A, Iñigo S, Fernández-Calvo P, Heyndrickx KS, Dhondt S, Shi H, De Milde L, Vanden Bossche R, De Clercq R, Eeckhout D, Ron M, Somers DE, Inzé D, Gevaert K, De Jaeger G, Vandepoele K, Pauwels L, Goossens A (2017) The transcriptional repressor complex FRS7-

Shiba Y, Takahashi T, Ohashi Y, Ueda M, Mimuro A, Sugimoto J, Noguchi Y, Igawa T (2023) Behavior of Male Gamete Fusogen GCS1/HAP2 and the Regulation in. Biomolecules 13:208

Shimada TL, Shimada T, Hara-Nishimura I (2010) A rapid and non-destructive screenable marker, FAST, for identifying transformed seeds of Arabidopsis thaliana. Plant J 61:519–528

Stuttmann J, Barthel K, Martin P, Ordon J, Erickson JL, Herr R, Ferik F, Kretschmer C, Berner T, Keilwagen J, Marillonnet S, Bonas U (2021) Highly efficient multiplex editing: one-shot generation of 8× Nicotiana benthamiana and 12× Arabidopsis mutants. Plant J 106:8–22

Swinnen G, De Meyer M, Pollier J, Molina-Hidalgo FJ, Ceulemans E, Venegas-Molina J, De Milde L, Fernández-Calvo P, Ron M, Pauwels L, Goossens A (2022) The basic helix-loop-helix transcription factors MYC1 and MYC2 have a dual role in the regulation of constitutive and stress-inducible specialized metabolism in tomato. New Phytol 236:911–928

Takahashi T, Mori T, Ueda K, Yamada P, Nagahara S, Higashiyama T, Sawada H, Igawa T (2018) The male gamete membrane protein DMP9/DAU2 is required for double fertilization in flowering plants. Development 145:dev170076

Tsutsui H, Higashiyama T (2017) pKAMA-ITACHI Vectors for Highly Efficient CRISPR/Cas9-Mediated Gene Knockout in Arabidopsis thaliana. Plant Cell Physiol 58:46–56

von Besser K, Frank AC, Johnson MA, Preuss D (2006) Arabidopsis HAP2 (GCS1) is a sperm-specific gene required for pollen tube guidance and fertilization. Development 133:4761–4769

Wang W, Xiong H, Zhao P, Peng X, Sun MX (2022) DMP8 and 9 regulate HAP2/GCS1 trafficking for the timely acquisition of sperm fusion competence. Proc Natl Acad Sci U S A 119:e2207608119

Weijers D, Franke-van Dijk M, Vencken RJ, Quint A, Hooykaas P, Offringa R (2001) An Arabidopsis Minute-like phenotype caused by a semi-dominant mutation in a RIBOSOMAL PROTEIN S5 gene. Development 128:4289–4299 positively charged carboxy-terminal domain. PLoS Genet 6:e1000882

