## Supplemental Table 1 for "Production of homozygous deletion mutants targeting fertilization regulator genes through multiplex genome editing"

**Supplementary Table 1 Primers used in this study.**

| Primer name | Sequence (5'-3') | Application |
|---|---|---|
| GCS1del5'g1FP | ATTGAAAAAACCCAAAGTAACCGT | Plasmid construction for *GCS1* deletion |
| GCS1del5'g1RP | AAACACGGTTACTTTGGGTTTTTT | Plasmid construction for *GCS1* deletion |
| GCS1del3'g1FP | ATTGGTAGTAGTGAGAGTCGCTG | Plasmid construction for *GCS1* deletion |
| GCS1del3'g1RP | AAACCAGCGACTCTCACTACTAC | Plasmid construction for *GCS1* deletion |
| GCS1del5'g2FP | ATTGAGTTGTGGTATACATTCCGT | Plasmid construction for *GCS1* deletion |
| GCS1del5'g2RP | AAACACGGAATGTATACCACAACT | Plasmid construction for *GCS1* deletion |
| GCS1del3'g2FP | ATTGATGTGCACGTGGAAAGACAG | Plasmid construction for *GCS1* deletion |
| GCS1del3'g2RP | AAACCTGTCTTTCCACGTGCACAT | Plasmid construction for *GCS1* deletion |
| GEX2del5'g1FP | ATTGGCCAACTGCTGTGATGTTTG | Plasmid construction for *GEX2* deletion |
| GEX2del5'g1RP | AAACCAAACATCACAGCAGTTGGC | Plasmid construction for *GEX2* deletion |
| GEX2del3'g1FP | ATTGAGCTTAACCAACTTAGATGA | Plasmid construction for *GEX2* deletion |
| GEX2del3'g1RP | AAACTCATCTAAGTTGGTTAAGCT | Plasmid construction for *GEX2* deletion |
| GEX2del5'g2FP | ATTGCTACATGCAGTGCACATCAG | Plasmid construction for *GEX2* deletion |
| GEX2del5'g2RP | AAACCTGATGTGCACTGCATGTAG | Plasmid construction for *GEX2* deletion |
| GEX2del3'g2FP | ATTGAGGTTGTGTATATATGTGTA | Plasmid construction for *GEX2* deletion |
| GEX2del3'g2RP | AAACTACACATATATACACAACCT | Plasmid construction for *GEX2* deletion |
| DMP8del5'g1FP | ATTGTCGAACTATCTTTAGTACGA | Plasmid construction for *DMP8* deletion |
| DMP8del5'g1RP | AAACTCGTACTAAAGATAGTTCGA | Plasmid construction for *DMP8* deletion |
| DMP8del3'g1FP | ATTGAAACGATGGATAGTAGTCAG | Plasmid construction for *DMP8* deletion |
| DMP8del3'g1RP | AAACCTGACTACTATCCATCGTTT | Plasmid construction for *DMP8* deletion |
| DMP8del5'g2FP | ATTGTGAATAAGTGAATATTTGCA | Plasmid construction for *DMP8* deletion |

| | | |
|---|---|---|
| DMP8del5'g2RP | AAACTGCAAATATTCACTTATTCA | Plasmid construction for *DMP8* deletion |
| DMP8del3'g2FP | ATTGTGTCCTACAATATAGTAGGT | Plasmid construction for *DMP8* deletion |
| DMP8del3'g2RP | AAACACCTACTATATTGTAGGACA | Plasmid construction for *DMP8* deletion |
| DMP9del5'g1FP | ATTGCTCTCTGATAACAAGCCACG | Plasmid construction for *DMP9* deletion |
| DMP9del5'g1RP | AAACCGTGGCTTGTTATCAGAGAG | Plasmid construction for *DMP9* deletion |
| DMP9del3'g1FP | ATTGTTGAGTCCGTGATGTAGTGG | Plasmid construction for *DMP9* deletion |
| DMP9del3'g1RP | AAACCCACTACATCACGGACTCAA | Plasmid construction for *DMP9* deletion |
| DMP9del5'g2FP | ATTGTATGGTTTGTCCGTAAATGA | Plasmid construction for *DMP9* deletion |
| DMP9del5'g2RP | AAACTCATTTACGGACAAACCATA | Plasmid construction for *DMP9* deletion |
| DMP9del3'g2FP | ATTGTGAACACTTATACAACTCAG | Plasmid construction for *DMP9* deletion |
| DMP9del3'g2RP | AAACCTGAGTTGTATAAGTGTTCA | Plasmid construction for *DMP9* deletion |
| GGfragment1_4FP | TTGGTCTCTAAGCCTTTTTTTCTTCTTCTTCGTT | Plasmid construction |
| GGfragment1_4RP | TTGGTCTCTAGTAGTCTAGAAAAAAAGCACCGAC | Plasmid construction |
| GGfragment2_4FP | TTGGTCTCTTACTCTTTTTTTCTTCTTCTTCGTT | Plasmid construction |
| GGfragment2_4RP | TTGGTCTCTCATTGTCTAGAAAAAAAGCACCGAC | Plasmid construction |
| GGfragment3_4FP | TTGGTCTCTAATGCTTTTTTTCTTCTTCTTCGTT | Plasmid construction |
| GGfragment3_4RP | TTGGTCTCTACCTGTCTAGAAAAAAAGCACCGAC | Plasmid construction |
| GGfragment4_4FP | TTGGTCTCTAGGTCTTTTTTTCTTCTTCTTCGTT | Plasmid construction |
| GGfragment4_4RP | TTGGTCTCTTTGCGTCTAGAAAAAAAGCACCGAC | Plasmid construction |
| gRNAsGGf1_2FP | TTGGTCTCTAAGCCTCGACGTCGCATGCCTGCAG | Plasmid construction for *DMP8/9* deletion |
| gRNAsGGf1_2RP | TTGGTCTCTAGTATTGAGCTCCAAGCTTGAATTC | Plasmid construction for *DMP8/9* deletion |
| gRNAsGGf2_2FP | TTGGTCTCTTACTCTCGACGTCGCATGCCTGCAG | Plasmid construction for *DMP8/9* deletion |
| gRNAsGGf2_2RP | TTGGTCTCTTTGCTTGAGCTCCAAGCTTGAATTC | Plasmid construction for *DMP8/9* deletion |

| | | |
|---|---|---|
| GCS1 del5P | ACGATGAGGAGCGTATTTCTCTTT | *GCS1* deletion check for T1-3 lines, sequencing |
| GCS1 del3P | TGAGTCTTTCTTTTCCTTCTGCTC | *GCS1* deletion check for T1-3 lines |
| GEX2 del5P | GCGGTTTGAAAAGTTCAAAA | *GEX2* deletion check for T1-3 lines. sequencing |
| GEX2 del3P | TAAACAAAGTCAATCTAATGCAGC | *GEX2* deletion check for T1-3 lines |
| GEX2 8ex5P | TAGTGTGACTCAATTCAATGG | *GEX2* deletion check for T1 lines |
| GEX2 ORF5P | AAGGAGATATCCATGGCGATTAAATTCGTTTCA | RT-PCR |
| GEX2 RT3P | TGGTCCTGAATTCACCTCAA | RT-PCR |
| GCS1 RT5P | TTCCCAGTGGATCCAGTGGAG | RT-PCR |
| GCS1 RT3P | CCGCCTCTCTTGGGATTACAAGAT | RT-PCR |
| DMP8 del5P | AAGACCAAGCAAAGGTTCAGT | *DMP8* deletion check for T1-3 lines, sequencing |
| DMP8 del3P | CGGAGTTGTAAATGAGAGGATAGT | *DMP8* deletion check for T1-3 lines |
| DMP9 del5P | TTAATTCTCTTGCGAAGGACTCT | *DMP9* deletion check for T1-3 lines |
| DMP9 del3P | TATCACATGACTTGAACATTCTCG | *DMP9* deletion check for T1-3 lines |
| DMP9delSEQ5P | CTTAAGTAATGTTTCATTTGATAG | Sequencing for *DMP9* deletion check |
| DMP9delSEQ3P | CTTGAATGAACTATATTGTAAAAG | Sequencing for *DMP9* deletion check |
