## Supplemental Figures for "Production of homozygous deletion mutants targeting fertilization regulator genes through multiplex genome editing"

(a)

| Gene | DeepSpCas9 score | | | | Deletion frequency (%) | Homozygous deletion acquisition rate (%) |
|---|---|---|---|---|---|---|
| | 5’g1 | 5’g2 | 3’g1 | 3’g2 | | |
| *GCS1* | 58.2 | 62.6 | 65.6 | 57.4 | 75.0 | 30.0 |
| *GEX2* | 44.5 | 61.3 | 52.1 | 62.0 | 40.0 | 20.0 |
| *DMP8* | 43.5 | 45.3 | 57.4 | 56.3 | 54.2 | 8.3 |
| *DMP9* | 65.8 | 59.8 | 54.8 | 70.2 | 63.6 | 18.2 |

(b)

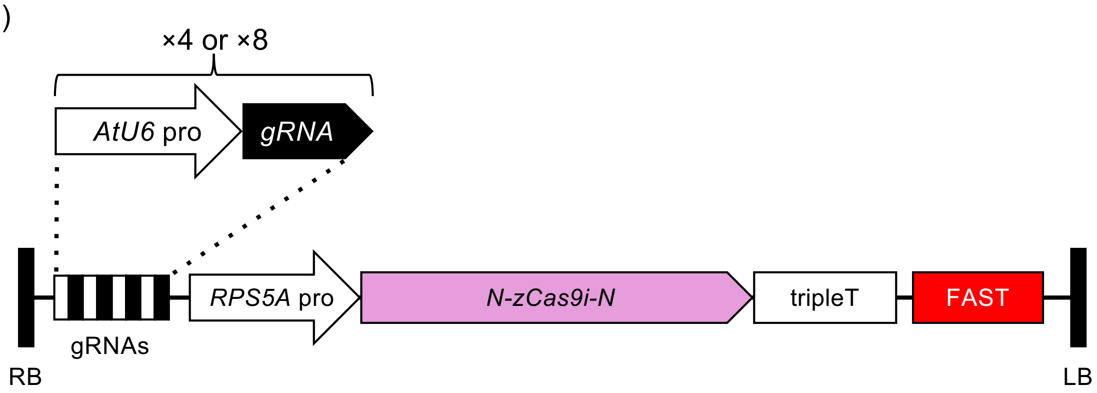


**Fig. S1** The scheme for gene deletion performed in this study.

**(a)** DeepSpCas9-predicted scores of the target sequences used for gene deletions. The percentages of gene deletion and homozygous deletion were calculated based on the genomic PCR results from each T1 generation (Figures 1a, 2a, and 3a). **(b)** T-DNA design used in this study. Four or eight guide RNA scaffold expression cassettes were inserted in the ‘gRNAs’ region. AtU6 pro; *A. thaliana* U6 gene promoter, RPS5A pro; *A. thaliana* ribosome protein subunit 5A gene promoter, *N-zCas9i-N*; *Zea mays*-codon optimized and intronized *Cas9* with nuclear localization signal sequences at both terminal ends. triple T; fusions of 35S terminator from Cauliflower Mosaic Virus, *Nicotiana benthamiana Actin3* terminator region, and the Rb7 matrix attachment region from *N. tabacum*. FAST; an expression cassette of a fusion gene of *A. thaliana Oleosin 1* and tagRFP (Shimada et al. 2010), RB; right border, LB; left border.

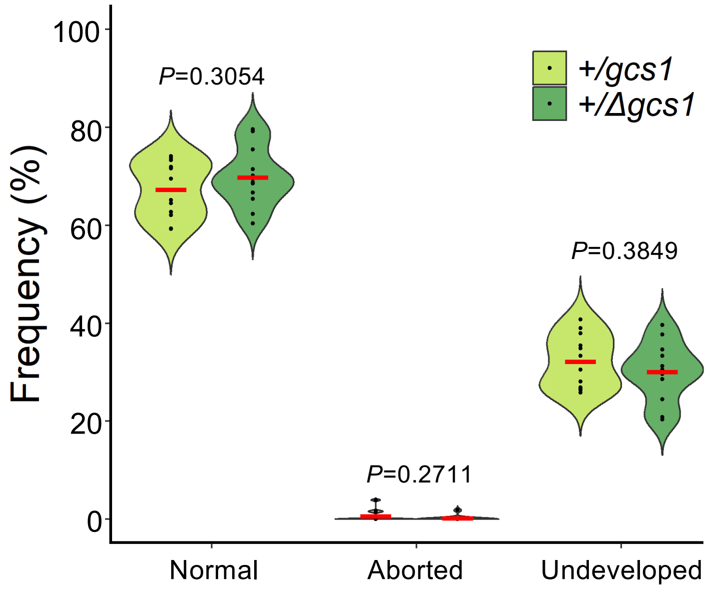


**Fig. S2** Comparison of the frequencies in each seed development phenotype between *gcs1* and *Δgcs1*.

Twelve siliques from each *gcs1* and *Δgcs1* heterozygous plant were dissected for seed counting. The frequencies of normal, aborted, and undeveloped seed phenotypes are represented with violin plots, in which +/*gcs1* and +/*Δgcs1* are colored in lime green and bright green, respectively. The black dots and a red horizontal bar in each plot represent each frequency per silique and the median. The P-values obtained by Welch's t-test are indicated above the plots.

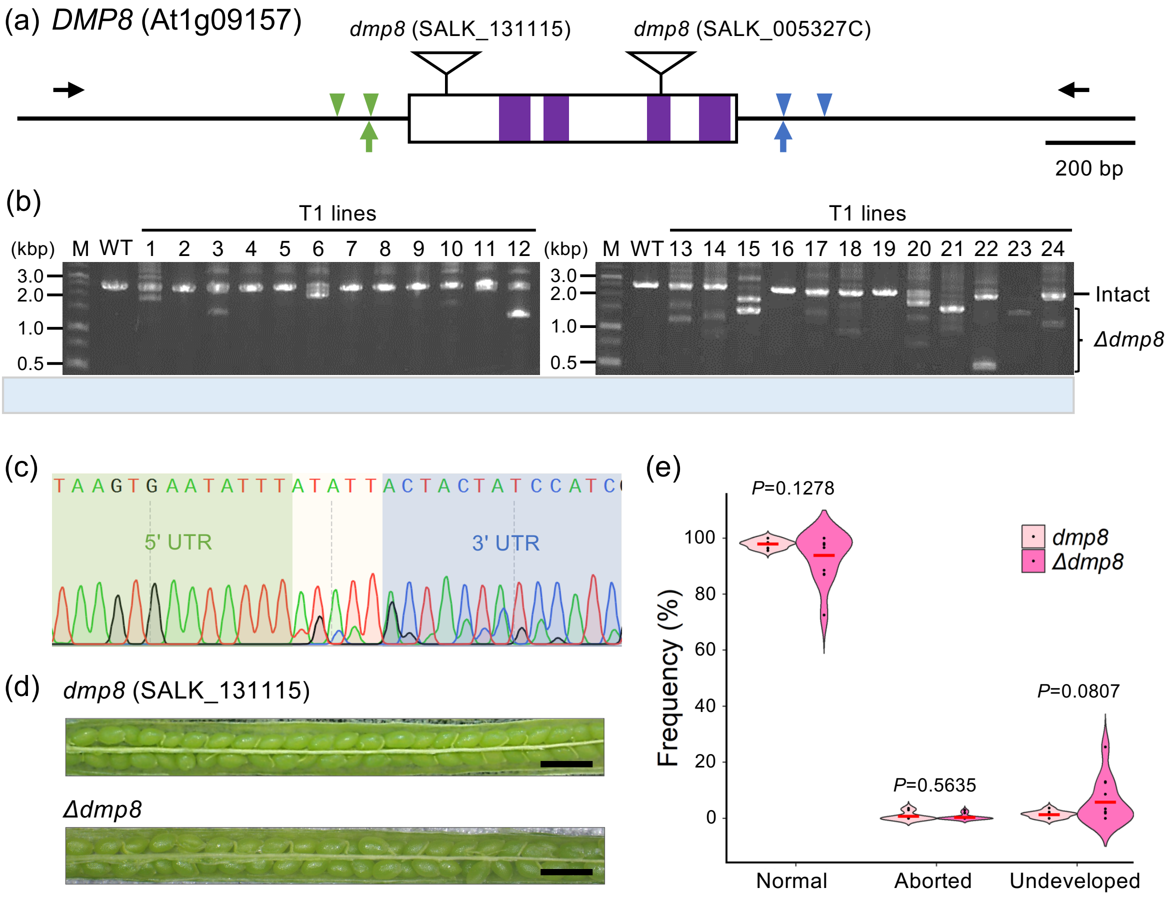


**Fig. S3** Gene deletion targeting *DMP8*

**(a)** Schematic drawing of genomic DMP9, an intronless gene. Green and blue triangles represent target sites at 5′ upstream and 3′ downstream of the ORF. Black arrows are primers used for genomic PCR to check the ORF deletion. Open triangles with a vertical line indicate the T-DNA insertion sites of the *dmp8* mutants used in the previous studies (SALK_13115 (Shiba and Takahashi et al. 2023); SALK_005327C (Wang et al. 2022). The corresponding transmembrane (TM) region is marked in purple. **(b)** Genomic PCR with primers shown in (a) to check the gene deletion in T1 transgenic lines. The positions of intact *DMP8* (ca. 2.5 kbp) and the gene deletion (*Δdmp8*) (ca. 1.1–1.5 kbp) amplicons are marked with a line and a curly bracket, respectively. The estimated zygosities of gene deletion based on the amplicon patterns are represented below the photo (H; homozygous, He; heterozygous, N; no deletion). **(c)** Sequencing result of a band from line 21 showing *DMP8* homozygous deletion (*Δdmp8*). **(d)** Siliques from *dmp8* (SALK_13115) and *Δdmp8* (lower). bar = 1 mm. **(e)** Violin plots of frequencies of each seed phenotype observed in *dmp8* (SALK_13115) (pale pink) and *Δdmp8* (pink) siliques. Nine and 12 siliques from *dmp8* and *Δdmp8* were dissected for seed counting. The black dots and a red horizontal bar in each plot represent each frequency per silique and the median. P-values between *dmp8* and *Δdmp8* detected with Welch's t-test are indicated above the plots.

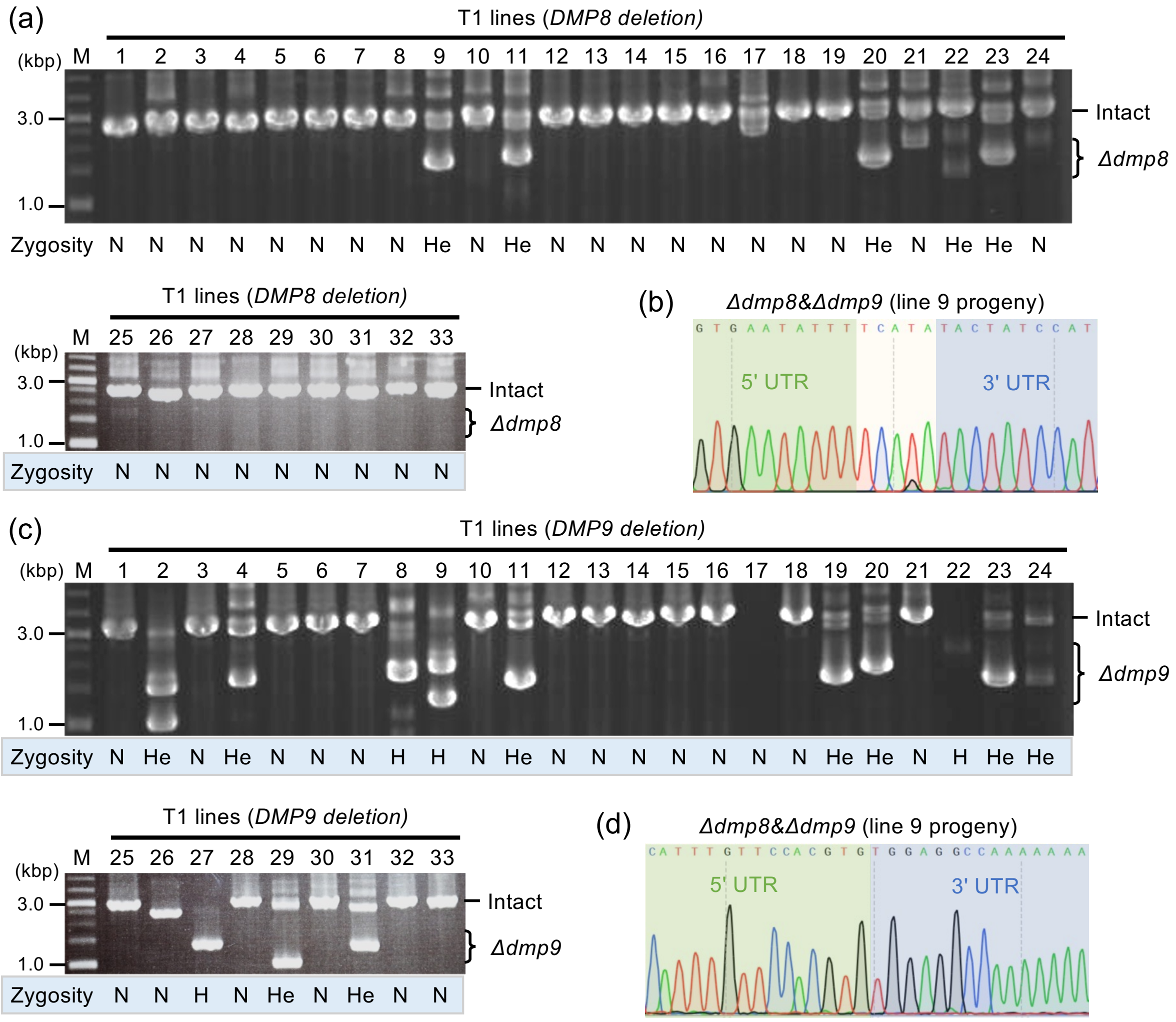


(e)

| Genotype | DMP8 DMP9 | DMP8 +/dmp9 | DMP8 dmp9 | +/dmp8 DMP9 | +/dmp8 +/dmp9 | +/dmp8 dmp9 | dmp8 DMP9 | dmp8 +/dmp9 | dmp8 dmp9 |
|---|---|---|---|---|---|---|---|---|---|
| The number of T1 lines | 20 | 6 | 2 | 0 | 3 | 2 | 0 | 0 | 0 |

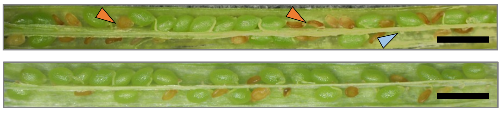


**Fig. S4** Simultaneous gene deletion targeting *DMP8* and *DMP9*

**(a)** Evaluation of *DMP8* deletion by genomic PCR with primers shown in Fig. S4(a) in the T1 transgenic lines. The positions of intact *DMP8* (ca. 2.5 kbp) and the gene deletion (ca. 1.1–1.5 kbp) amplicons are marked with a line and a curly bracket, respectively. The estimated zygosities of gene deletion based on the amplicon patterns are represented below the photo (H; homozygous, He; heterozygous, N; no deletion). **(b)** Verification of *DMP8* homozygous deletion in the line 9 progeny by sequencing. **(c)** Evaluation of *DMP9* deletion by genomic PCR with primers shown in Fig. 3(a) in the same T1 transgenic lines in (a). The intact *DMP9* amplicon (ca. 2.9 kbp) and the smaller bands suggesting deletions (ca. 1.0–1.7 kbp) are marked with a line and a curly bracket, respectively. The estimated zygosities of gene deletion based on the amplicon patterns are represented below the photo as in (a). M; size marker. **(d)** Verification of *DMP9* homozygous deletion in the line 9 progeny by sequencing. **(f)** Number of obtained T1 lines with various *DMP8* and *DMP9* deletion genotypes. **(g)** Siliques from *dmp8/9* (*dmp8* (SALK_13115) and *dmp9*$^{GE}$ double mutant) and *Δdmp8&Δdmp9* obtained in this study. Pale blue and orange triangles indicate aborted and undeveloped seeds. bar = 1 mm.
